# MS4MS: LLMs-driven Multi-agent System for Small-molecule Identification via LC-MS/MS

**DOI:** 10.64898/2025.12.02.691830

**Authors:** Na Guo, Jianbin Guo, Yizhe Liu, Sibo Wei, LiFeng Dong, Hongmin Du, Yang Bai, Yurou Zhao, Xiaoqing Wang, Dajun Zeng, Hongjun Yang

**Affiliations:** Experimental Research Center, China Academy of Chinese Medical Sciences, Beijing, 100700, China; Beijing Key Laboratory of Digital-Intelligent Equipment of Traditional Chinese Medicine Four Diagnostic Techniques, Beijing, 100041, China; Institute of Automation, Chinese Academy of Sciences, Beijing, 100190, China; Beijing Wenge Technology Co.,Ltd, Beijing, 100190, China; School of New Media and Communication, Tianjin University, Tianjin, 300350, China; School of Pharmacy, Shenyang Pharmaceutical University, Shenyang, 110016, China

## Abstract

Small molecule identification is central to research fields such as drug discovery, but in complex systems like Traditional Chinese Medicine (TCM), traditional mass spectrometry analysis methods remain constrained by bottlenecks including insufficient database coverage, fragmented analysis workflows, and poor result interpretability. To address these limitations, we developed MS4MS, a large language model-driven multi-agent system that enables an end-to-end automated pipeline from raw data to small molecule identification. Validation on a public benchmark demonstrates that MS4MS achieves state-of-the-art performance in molecular formula prediction. Furthermore, its innovative small molecule identification agent enables efficient and interpretable compound elucidation. Verification using herbal extracts indicates MS4MS’s outstanding performance regarding analytical coverage and the discrimination of isomers. Consequently, MS4MS offers a novel, accurate, interpretable, and high-throughput end-to-end automated strategy for small molecule identification, overcoming the analytical bottlenecks of traditional mass spectrometry in natural products and complex TCM systems.

## Introduction

Small molecules, including metabolites, natural products, drugs, and their degradation products, are core elements constituting the chemical basis of life and play a crucial role in fields such as drug discovery^1^. Over the last several decades, rapid advancement in analytical chemistry, particularly the enhanced capabilities of liquid chromatography (LC) combined with tandem mass spectrometry (MS/MS)^2^, have significantly improved our ability to detect these chemical entities^3^. However, an increasingly widening gap has emerged between the rapid advancement of data acquisition capabilities and the sluggish improvement in data interpretation efficiency. Transforming massive volumes of mass spectrometry data into reliable chemical structure information, namely small molecule identification, remains a central bottleneck in the field. The root of this bottleneck lies in the inherent complexity of small molecule fragmentation mechanisms. In contrast to macro molecules like proteins or DNA that feature linear structures and relatively clear rules, small molecules exhibit highly varied non-linear architectures^4^. Their fragmentation behaviors in mass spectrometry are intricately influenced by atomic connectivity, stereochemistry, localized charge distribution, and collision energy, making spectrum prediction and interpretation a knowledge-intensive process that relies heavily on expert experience and intuitive judgment. This challenge is particularly acute in the traditional Chinese Medicine (TCM) small molecule identifications^5–10^. As a prototypical complex chemical system, a TCM formula derives its pharmacological effects from the synergistic action of multiple constituents. Natural products display extremely high structural diversity and complexity, and the presence of numerous isomers and trace-level metabolites further complicates the analytical landscape. Faced with those “chemical dark matter”, conventional identification methods often fall short, creating a severe bottleneck that limits our ability to comprehensively and systematically elucidate the pharmacologically active substances in TCM and to leverage their potential for modern drug discovery.

The emergence of artificial intelligence for science (AI4S), exemplified by breakthroughs such as AlphaFold^11^ in protein structure prediction, has catalyzed a paradigm shift across multiple scientific domains^12,13^. In metabolomics, this offers new opportunities to overcome long-standing limitations in small molecule identification^14,15^. The integration of AI with high-resolution mass spectrometry enables the potential for high-accuracy, scalable annotation of complex natural product systems^16,17^.

The application of AI in small molecule identification is currently at a critical juncture of methodological innovation. Molecular formula prediction is the starting point for small molecule identification, primarily divided into two steps: candidate chemical formula generation and unique molecular formula determination^18^. During the candidate formula generation stage, prevalent approaches typically depend on brute-force algorithms that constrain the elemental composition and counts^19,20^. Such algorithms not only require manual setting of elemental constraints, but have high complexity, often resulting in either missing the correct formula or including a large number of false positives. Another category of methods, based on bottom-up MS/MS interrogation^21^, is heavily dependent on the quality of pre-built databases and is susceptible to interference from spectral noise peaks. The subsequent isomer identification step also faces multiple challenges. Rule-based methods are constrained by the scalability of handcrafted rules and difficulties in balancing weights^22–24^, while methods based on machine learning or deep neural network, despite showing high accuracy on specific datasets, generally suffer from poor transferability and interpretability^20,21,25,26^. For small molecule identification, the most common method is spectral library matching^22,27,28^. These methods relies heavily on database coverage and the robustness of scoring algorithms^29,30^, and is ineffective for “dark matter” not present in the library. To reduce the reliance on databases, in-silico fragmentation simulation and scoring techniques emerged^31^. Nevertheless, while such methods can generate extensive fragmentation data, the credibility of the simulated outcomes is constrained, making them an inadequate substitute for real experimental data. Concurrently, these methods are also limited by issues such as low coverage of fragmentation rules^19^, poor interpretability^16^, or difficulty in balancing scoring weights^32^. To avoid the biases inherent in silico fragmentation, some researcher focus on de novo molecular prediction^33–35^. This method does not interpret MS/MS fragment peaks but instead directly predicts molecular fingerprints or structures, leading to a decision-making process that lacks chemical intuition or biosynthetic background, thereby limiting the application of the identification results in scientific research.

To address the above challenges, we propose MS4MS, a multi-agent system based on large language models^36^, designed to achieve full-workflow small molecule identification. MS4MS implements an end-to-end automated process from raw spectral data parsing to small molecule identification, organized around a spectrum processor, a molecular formula prediction agent, a small molecule identification agent, and a reporting agent. In contrast to previous methods, our molecular formula prediction agent achieves efficient and accurate molecular formula prediction without the need for manual constraints. Furthermore, the small molecule identification agent revolutionizes the small molecule identification paradigm, implementing an accurate and high-throughput, and fragmentation-mechanism interpretable identification. More importantly, it is capable of elucidating semi-unknown and completely unknown compounds that are structurally similar to known small molecules, thereby expanding the research boundaries of small molecule identification. To evaluate the effectiveness of MS4MS, we conducted experiments on public benchmark datasets, and the results show that our method achieves state-of-the-art performance in molecular formula prediction. Meanwhile, to verify the efficacy of the system, we applied it to the analysis of three representative TCM compound formula: Buyang Huanwu decoction, Dengzhan Shengmai granule, and Zhuodu Huanwu decoction. These formulas feature complex compositions covering multiple structural types such as flavonoids, saponins, alkaloids, and phenolic acids, providing an ideal test platform to evaluate the agent’s identification capability, isomer differentiation ability, and potential for discovering new structures in real-world complex scenarios. This study not only provides a novel and reliable automated solution for small molecule identification but also lays a methodological foundation for elucidating the chemical basis of complex systems like TCM and accelerating their application in modern drug development. MS4MS has been integrated into our online platform, which researchers can directly access and use.

## Results

### Problem definition

Given an LC-MS/MS dataset 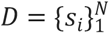 in mzML format, where each spectrum *s*_*i*_ = (*MS*1_*i*_, *MS*2_*i*_, *meta*_*i*_) contains precursor ion information, fragment peaks, and associated metadata (e.g. retention time), the goal is to identify the most probable molecular structure *M*^∗^ corresponding to each spectrum.

#### Step 1. Spectrum Parse

From each MS1 spectrum, extract the precursor ion’s m/z value ( *m*/*z*_*p*_), charge ( *z*) and adduct type ( *A*). From the corresponding MS2 spectrum, collect the set of fragment ion peaks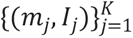, where *m*_*j*_ is the m/z of the j-th fragment ion, and *I*_*j*_ is its relative intensity. This yields:

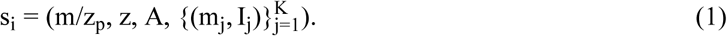

#### Step 2. Molecular Formula Inference

Based on the precursor *m*/*z*_*p*_, isotopic distribution, adduct type, and fragmentation information from MS2, the most plausible molecular formula *F*^∗^ is inferred as:

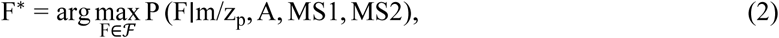

where ℱ is the set of chemically valid formulas constrained by elemental and isotopic rules. MS2 fragment ions help validate or discard candidate formulas that are inconsistent with the observed fragmentation pattern.

#### Step 3. Small Molecule Identification

Given the inferred molecular formula *F*^∗^, identify the most probable structural candidate *M*^∗^ from a database of compounds by matching its predicted or experimental fragmentation pattern:

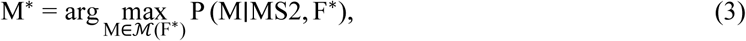

where ℳ(*F*^∗^) is the set of all structural isomers consistent with formula *F*^∗^, and the probability term *P*(*M*∣MS2, *F*^∗^) reflects the likelihood that observed MS2 fragmentation originates from structure *M*.

### LLM-based multi-agent system for small molecule identification

We developed and validated MS4MS, a novel large language models (LLMs)-based multi-agent system, created to tackle the existing problems of accuracy, efficiency, and process interpretability in small molecule identification. MS4MS pioneeringly integrates an end-to-end automated pipeline, from raw mass spectrometry data analysis to high-confidence compound structure identification. The overall architecture of MS4MS is illustrated in Fig. 1. It is constructed around two core analytical agents, the molecular formula prediction agent and the small molecule identification agent, and builds a rigorous, transparent, and efficient analytical pipeline in collaboration with a spectrum processor and a reporting agent.

**Fig. 1.**
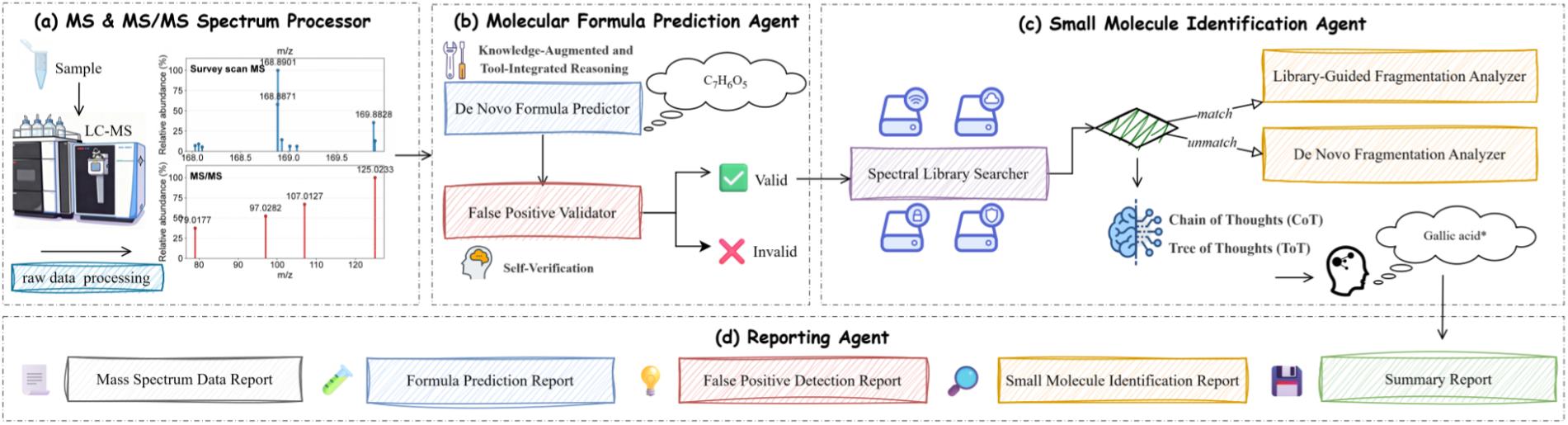
Architecture of MS4MS

The identification process commences with high-accuracy molecular formula prediction. Traditional methods at this stage are typically constrained by predefined elemental sets and their quantities, significant algorithmic complexity, or a critical dependency on high-quality MS/MS data and external databases^20,21,25,26^. In contrast, our molecular formula prediction agent overcomes these deficiencies by leveraging the vast chemical knowledge internalized by the LLM to achieve intelligent molecular formula prediction without requiring manually set elemental constraints. This agent can automatically skips chemically unreasonable combinations, thereby substantially trimming the candidate molecular formula space and enabling efficient generation of candidate formulas. Meanwhile, to ensure reasoning accuracy, the agent also integrates tool based on the model context protocol (MCP)^37^ service to mitigate the risk of LLM hallucinations in precise molecular weight calculations, and employs a critical self-verification^38^ mechanism to cross-validation signal quality, effectively eliminating false positive interference. This architecture guarantees that only authentic, high-quality signals enter the downstream analysis, thereby boosting the efficiency and accuracy of the entire identification pipeline at its origin.

After obtaining a high-confidence molecular formula, the small molecule identification agent employs a dual-path analysis strategy to address identification challenges under different scenarios. Initially, the agent conducts a similarity retrieval of the experimental spectrum against several spectral libraries. If a matching spectrum is found, it will activate the library-guided fragmentation analyzer for in-depth validation. The core advantage of the library-guided fragmentation analyzer is that it provides interpretability of the fragmentation mechanisms. Traditional spectral library matching typically only provides a black-box similarity score, failing to explain the chemical principles of the match or the existing discrepancies^27,28^. However, the library-guided fragmentation analyzer resolves this critical issue. It does not simply assign a rigid score but rather systematically aligns the experimental and library spectra, attempting to associate the primary observed fragment ions with chemically sound fragmentation pathways, thereby offering a secondary verification for high-scoring matches. More importantly, it can deeply analyze the subtle differences between the two spectra, identify which peaks are background noise, or explain that certain mass differences are caused by specific group modifications, thereby significantly increasing the credibility of the result.

One notable contribution of the small molecule identification agent is that it bridge the gap between the known and unknown chemical space. For the library-guided fragmentation analyzer, even when faced with an imperfectly matched, semi-unknown compound, it can still perform derivative analysis based on structural similarity, thus expanding the application boundaries of existing spectral libraries. When faced with a compound that has no match in the library at all, the de novo fragmentation analyzer provides powerful exploratory capabilities. On one hand, this analyzer can perform de novo analysis directly from the experimental spectrum to attempt to elucidate completely unknown compounds, and on the other hand, it can also utilize structurally similar compounds to conduct derivative analysis of semi-unknowns.

### Molecular formula prediction agent improve the accuracy of molecular formula prediction

As the crucial entry to the small molecule identification, the accuracy of molecular formula prediction directly determines the success or failure of downstream analysis. To quantitatively evaluate the performance of our proposed molecular formula prediction agent, we compared it against several state-of-the-art molecular formula prediction methods on a public benchmark dataset containing 17,556 spectral entries, including MIST-CF^26^, FIDDLE^25^, BUDDY^21^, and SIRIUS^20^.

The evaluation results are shown in Table 1. Our agent, BUDDY, and MIST-CF achieved a 100% prediction success rate, whereas FIDDLE, SIRIUS 6, and SIRIUS 6 + zodiac failed to make predictions on 8412, 109, and 142 samples, respectively, demonstrating the superior robustness of our method. Regarding the overall top-1 accuracy, our agent achieved 92.04%, ranking highest among all evaluated methods. In comparison, the automated tools BUDDY and SIRIUS only achieved scores of 47.89% and 58.63%, respectively. Meanwhile, the deep learning methods MIST-CF and FIDDLE performed poorly without fine-tuning, scoring only 36.36% and 8.87%, which exposes their weakness of poor transferability.

**Table 1.** Evaluation results of baseline methods and our method on the MassSpecGym test set.

| Method | Succeed Samples | Failed Samples | Top-1 Acc.<br>(Succeed) | Overall Top-1<br>Acc. |
| --- | --- | --- | --- | --- |
| MIST-CF | 17556 | 0 | 36.36% | 36.36% |
| FIDDLE | 9144 | 8412 | 17.03% | 8.87% |
| BUDDY | 17556 | 0 | 47.89% | 47.89% |
| SIRIUS 6 | 17447 | 109 | 49.39% | 49.08% |
| SIRIUS 6 + zodiac | 17414 | 142 | 59.11% | 58.63% |
| Ours | 17556 | 0 | **92.04%** | **92.04%** |

### Comprehensive compound identification from TCM formula and experimental validation

To construct a high-quality, reliable dataset for subsequent MS4MS evaluation, we systematically investigated three classic TCM formulae with demonstrated pharmacological activities and clinical value: Buyang Huanwu Decoction (BYHW), Dengzhan Shengmai granule (DZSM), and Zhuodu Huanwu Decoction (ZDHW) decoctions. These complex formulae contain diverse structural classes including flavonoids, saponins, alkaloids, and phenolic acids, alongside numerous isomers and trace constituents. Using a robust non-targeted liquid chromatography-tandem mass spectrometry (LC-MS/MS) approach, a substantial number of compounds were successfully identified.

Detailed identification results are provided in Table 5. We identified 180, 205, and 183 compounds from DZSM, BYHW, and ZDHW decoctions, respectively. Crucially, to ensure maximum reliability, a significant subset of these identifications was unambiguously confirmed using commercial reference standards analyzed under identical conditions. Specifically, 73 compounds in DZSM, 75 in BYHW, and 68 in ZDHW were verified through this method, establishing a “gold-standard” subset within our dataset. The remaining compounds were assigned with high confidence through integration of high-resolution accurate mass measurements, characteristic MS/MS fragmentation patterns, and cross-referencing with authoritative spectral libraries.

To demonstrate our rigorous validation approach, we selected representative key bioactive components from major chemical classes for detailed presentation. Fig. 2 shows representative identification examples. The excellent agreement between samples and reference standards in their primary mass spectra, MS/MS fragmentation patterns, and retention times provides definitive evidence for compound identification.

**Fig. 2.**
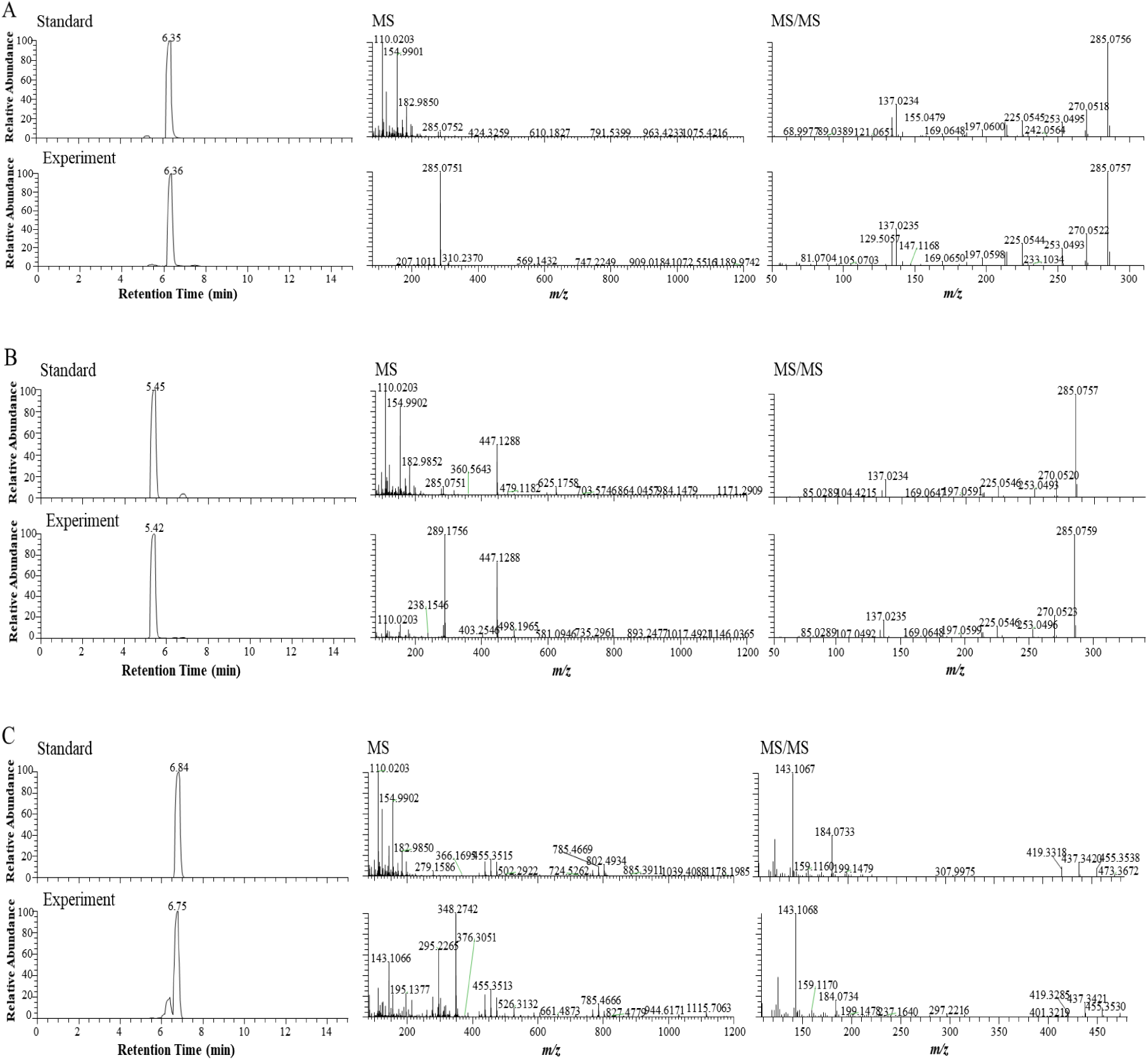
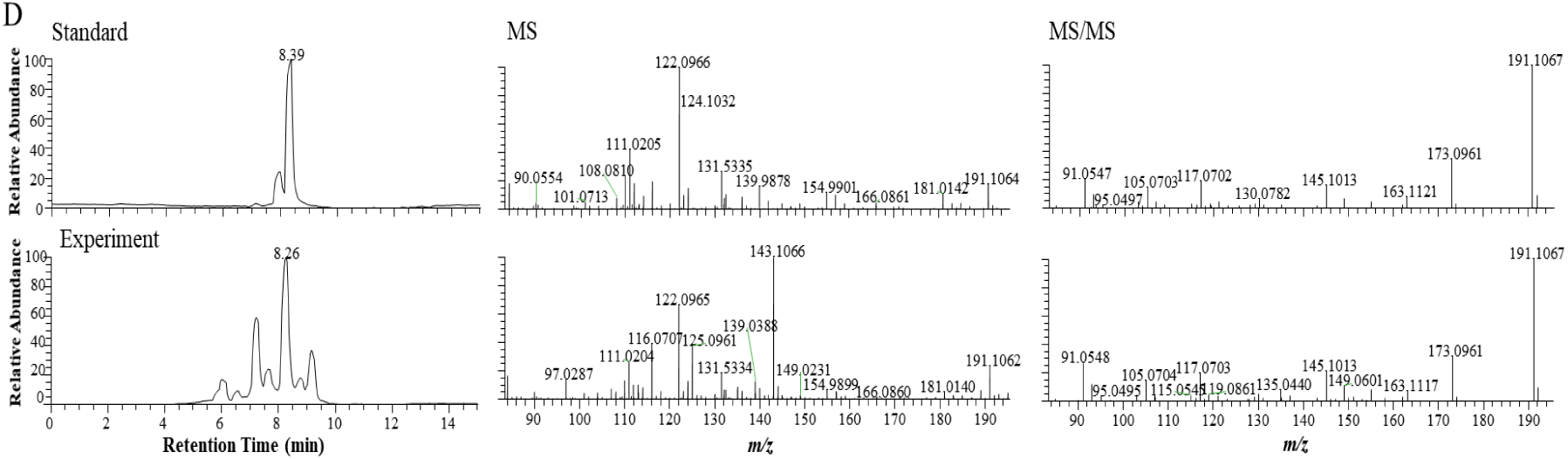
Experimental validation of major chemical constituents. (A) Calycosin. (B) Calycosin 7-O-Clucoside. (C) Astragaloside IV, (D) Ligustilide.

**Fig. 3.**
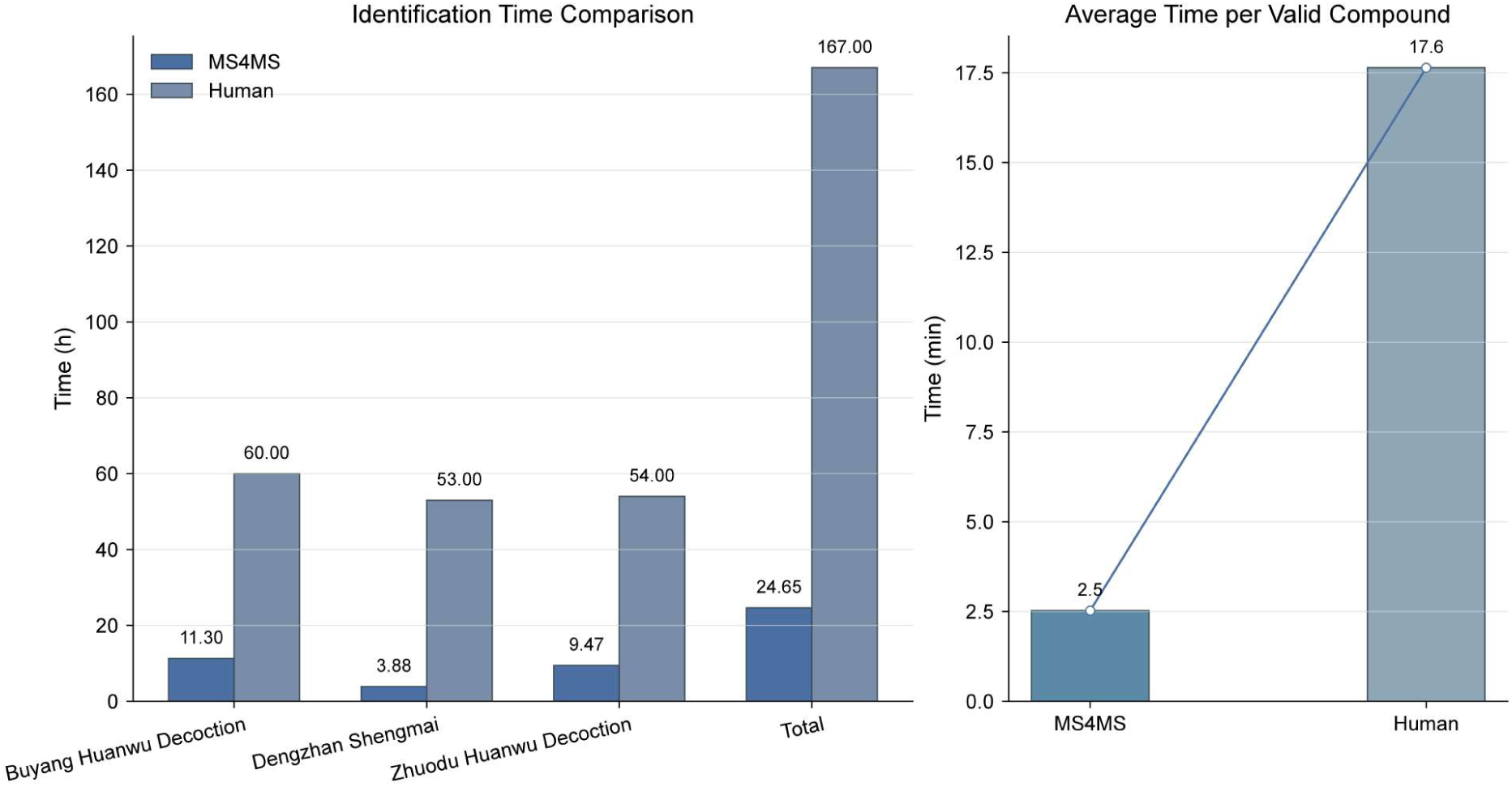
MS4MS vs Human identification time comparison

In summary, through meticulous wet laboratory experiments, we have established a comprehensive and highly reliable chemical inventory for three complex TCM formulae. This dataset, particularly the subset of compounds confirmed by reference standards, provides a valuable ground-truth benchmark. This solid experimental foundation is crucial for subsequent development and rigorous testing of MS4MS, ensuring its learning and predictions are based on verified data rather than speculative annotations.

Subsequently, we used the MS4MS system to perform a fully automated, end-to-end identification analysis on the same raw data. As shown in Table 2, MS4MS identified 211, 179, and 196 small molecules in the three formulas, respectively, achieving accuracies of 96.81%, 100%, and 92.98%, while maintaining identification numbers close to the manual results. This result indicates that MS4MS not only possesses high-throughput processing capabilities, but its systematic identification strategy also effectively reduces missed detections caused by human fatigue or subjective bias. To further quantify its reliability, we calculated the coverage of MS4MS identification results against the manual benchmark list. The results showed that its coverage for BYHW, DZSM, and ZDHW reached 88.12%, 93.14%, and 93.29%, respectively. This superior coverage performance proves that MS4MS can efficiently and accurately reproduce the vast majority of known components verified by humans, ensuring the biological relevance of the identification results.

**Table 2.** Experimental results of the end-to-end small molecule identification workflow using MS4MS. Coverage is defined as the ratio of the number of identifications in the intersection of MS4MS and Human results to “Identified (Human)”.

| Formula | Identified (MS4MS) | Accuracy | Identified (Human) | Coverage |
| --- | --- | --- | --- | --- |
| Buyang Huanwu Decoction (BYHW) | 188 | 96.81% | 202 | 88.12% |
| Dengzhan Shengmai Granule (DZSM) | 171 | 100% | 175 | 93.14% |
| Zhuodu Huanwu Decoction (ZDHW) | 171 | 92.98% | 164 | 93.29% |

More importantly, MS4MS demonstrated strong potential for new discoveries. As shown in Table 3, among all identified molecules, a portion were components unreported in the manual analysis. To validate the reliability of these novel annotations, we commissioned a third-party agency to conduct blind validation on MS4MS-unique annotations, confirming accuracy rates of 40%, 100%, and 45.45%, respectively.

**Table 3.**
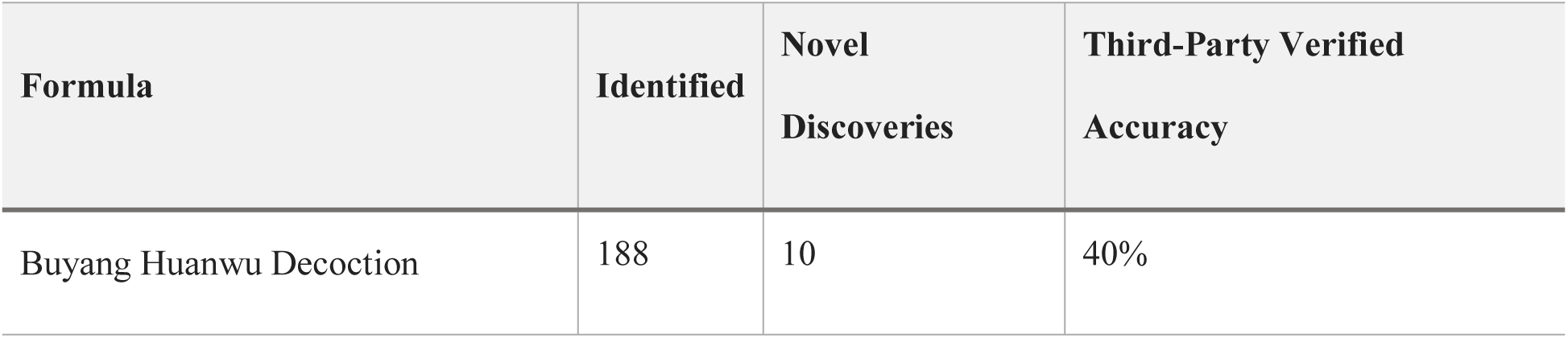
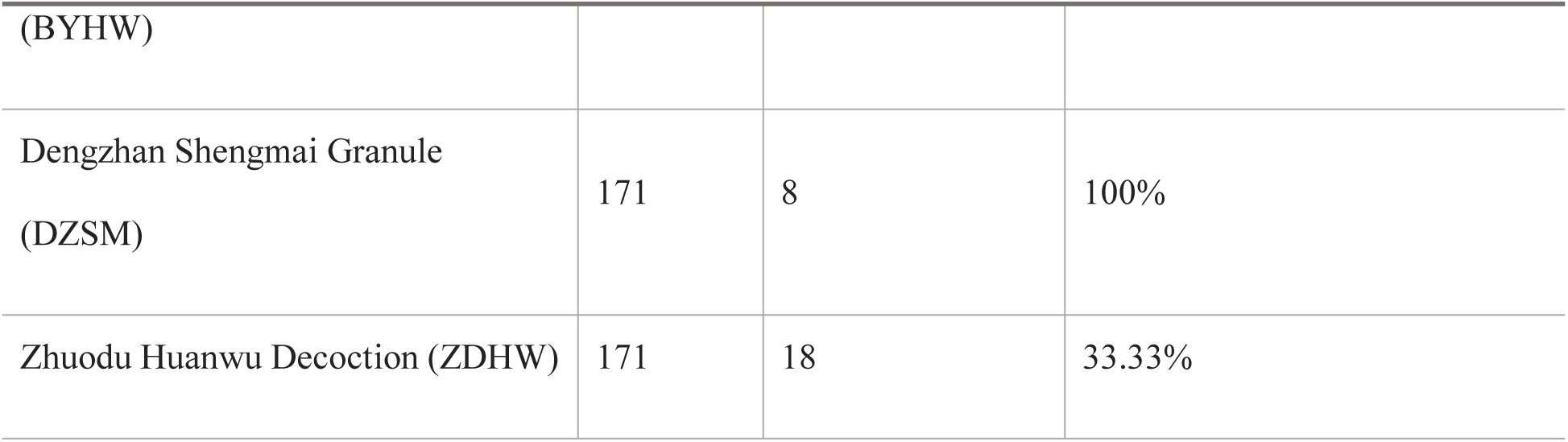
Third-party validation results of small molecules exclusively identified by MS4MS.

Moreover, regarding analysis efficiency, MS4MS demonstrated a significant advantage in high-throughput processing as an automated end-to-end tool. Illustrated in Fig. 4, in terms of total time, completing the analysis of the three formulas manually took 167 hours, while MS4MS required only 37.65 hours, representing a nearly 6.8-fold increase in overall efficiency. This high-throughput characteristic was even more prominent in terms of average efficiency, where MS4MS averaged just 2.5 minutes per valid compound identification, whereas manual identification averaged 17.6 minutes, marking a nearly 7-fold boost in unit efficiency.

## Discussion

In this paper, we present an LLM-based multi-agent system, MS4MS, which represents a paradigm shift. It is not just an automated tool, but rather an analytical system capable of reasoning. By integrating the reasoning power of large scientific foundation models with stringent chemical verification logic, we have achieved breakthroughs in the efficiency, accuracy, and procedural interpretability of small molecule identification.

We utilized the molecular formula prediction agent to achieve outstanding performance in molecular formula prediction, which establishes a robust cornerstone for the subsequent identification pipeline. As shown in Table 1, our agent achieved an overall top-1 accuracy of 92.04% on a large-scale test benchmark, significantly outperforming traditional methods. This is attributable to our agent avoiding several major limitations inherent in traditional methods. During the candidate molecular formula generation phase, traditional methods often depend on brute-force algorithms requiring manually defined elemental constraints and have high algorithmic complexity^19^. Even the bottom-up MS/MS interrogation approach is critically dependent on the quality of pre-built databases and is easily perturbed by noise peaks in low-quality MS/MS data^21^. In the small molecule identification stage, rule-based methods are limited by the scalability of manual rules, and the weights between rules are difficult to balance, often leading to trade-offs^19^. Methods based on traditional deep learning, on the other hand, suffer from poor interpretability, weak transferability, and a high barrier to entry requiring specialized modeling knowledge^26^. In contrast, our method does not require manually set elemental constraints, utilizing the vast chemical knowledge internalized by the LLM to automatically prune the chemically implausible candidate space. This avoids the invalid and potentially erroneous combinatorial explorations of traditional methods, and being independent of external databases, is more robust to variations in spectral quality. Critically, its reasoning process is flexible and interpretable, capable of dynamically adjusting analytical weights based on the specific situation rather than relying on rigid rules. This, combined with tool integration and a self-verification mechanism, ensures computational accuracy and the rejection of false positives, leading our method to its outstanding 92.04% performance.

Besides the absolute lead in overall top-1 accuracy, the failed samples in Table 1 further highlight the robustness and applicability of our method. Our agent, MIST-CF^26^ and BUDDY^21^, successfully predicted all 17,556 samples. By comparison, FIDDLE^25^ exhibited significant limitations, failing on 8,412 samples and thus reaching a coverage of less than 50%. Similarly, SIRIUS^20^ 6 and its zodiac^39^ variant also had 109 and 142 failed samples, respectively. This deficiency in coverage causes a discrepancy between the successfully predicted top-1 accuracy and the overall top-1 accuracy. In contrast, our method achieved a 100% prediction success rate, confirming that our approach is not only highly accurate but also possesses stronger applicability.

By successfully applying the MS4MS system to the identification of Buyang Huanwu decoction, Dengzhan Shengmai granule, and Zhuodu Huanwu decoction, this study fully validates its superior performance in handling the highly complex chemical systems of TCM. As demonstrated in Table 2, MS4MS outperformed manual analysis in terms of identification throughput, sustained a high average accuracy of 96.60%, and realized a excellent average coverage of 91.52% relative to manual benchmarks. Crucially, the system delivered a significant improvement in identification efficiency, evidenced by a nearly 6.8-fold reduction in total analysis time and an approximate 7-fold increase in the efficiency of single-compound identification.

The comprehensive advantages of MS4MS in identification breadth, accuracy, and efficiency stem from the fact that our designed small molecule identification agent addresses three major challenges in current identification methods. First, traditional spectral library matching methods cannot explain unmatched peaks or potential interferences, and their effectiveness is limited by the coverage of the spectral library^22,28^. Second, in-silico simulation methods face problems of poor interpretability and difficulty in balancing scoring weights^16,32^. Third is the helplessness when faced with unknown compounds. Compared to previous methods, our agent can not only answer “if it matches” but also explain “why it matches” and “why it is an imperfect match” through fragmentation mechanisms analysis. This capability is profoundly significant because it not only confirms the reliability of high-scoring matches with chemical principles but also reduces the probability of misidentification by traditional scoring algorithms due to instrumental noise or minor structural differences. More importantly, the agent possesses the capability to identify semi-unknown and even completely unknown compounds. This greatly expands the application boundaries of existing spectral libraries and provides a computational engine for exploring chemical “dark matter”.

Despite the progress described above, our method still possesses certain limitations. First, current LC-MS experiments and the subsequent small molecule identification are in a relatively independent state, which creates a fragmented workflow, preventing data analysis results from providing real-time feedback to guide experimental parameter optimization, thereby limiting further improvements in overall analytical efficiency and interpretation depth. Second, MS/MS data has inherent physicochemical limitations in distinguishing stereoisomers and certain regioisomers. Regardless of the reasoning power of LLMs, their judgment is always limited by the upper bound of information provided by one-dimensional spectra, making it difficult to overcome the common challenge of precise isomer differentiation. Finally, the professionalism and confident tone of the reports generated by the agent may inadvertently lead researchers to lower their critical review of its conclusions, introducing a risk of automation bias.

In the future, we have two primary research objectives. The first objective is to achieve embodied intelligence^40^, integrating the experimental instruments with the small molecule identification agent to realize an end-to-end, intelligent closed loop from sample analysis to data interpretation, enabling the agent to dynamically adjust instrument parameters based on real-time identification results. The second is to integrate multi-dimensional information. Future agents will need to perform reasoning in multi-dimensional space, not just rely on 1D data. This presents a challenge for efficient information representation, as compressing vast multi-dimensional data into a unified, compact, and low-loss embedding space will be essential for enabling efficient and accurate isomer discrimination. Following this developmental trajectory, we anticipate that agents will go beyond simply identifying a compound to reasoning about its origin (“where it comes from”) and transformation (“how it transforms”), thereby bridging the gap between mere identification and comprehensive understanding, and providing strong methodological support for research on complex TCM matrices and small molecule analysis.

In conclusion, MS4MS is not just a performance improvement over existing identification tools, but an AI-empowered paradigm shift. It opens up a new end-to-end automated and process-interpretable path for small molecule identification research, which will vigorously promote scientific exploration and discovery in this field.

## Methods

### Design and implementation of the MS4MS system

#### Architecture

MS4MS is functionally partitioned into four core components, which correspond to (a) the MS & MS/MS Spectrum Processor, (b) the Molecular Formula Prediction Agent, (c) the Small Molecule Identification Agent, and (e) the Reporting Agent, as illustrated in Fig. 1. The overall pipeline of the system is as follows:

1. Data Input and Preprocessing: The pipeline begins with the (a) MS & MS/MS Spectrum Processor. This processor is responsible for receiving and parsing the raw mzML format mass spectrometry data, and performing adduct ion deconvolution on the parsed data, thereby transforming the raw data into an LLM-comprehensible format.
2. Molecular Formula Prediction and Validation: The (b) Molecular Formula Prediction Agent receives a prompt composed of experimental spectral data from (a), is driven by the molecular formula prediction model, and utilizes knowledge augmentation and tool integration to enhance reasoning performance. Knowledge augmentation is realized via the extensive, implicit chemical knowledge that the foundation model internalizes during its pre-training^41^ and post-training^42^ phases. Such knowledge encompasses rules of chemical stability, the composition of common functional groups, atomic valence constraints, and isotopic abundance patterns, among others. This embedded knowledge functions as a robust heuristic filter, ensuring the model avoids exploring chemically unstable or implausible combinations during inference, thereby drastically pruning the vast candidate molecular formula space. Tool integration is implemented via the MCP^37^ service to mitigate the risk of model hallucinations, particularly in aspects like floating-point calculations. The model will perform tool calls within the corresponding service when molecular weight calculations are involved, ensuring the accuracy and reliability of the computations. Furthermore, the agent implements a self-verification^38^ mechanism, the core logic of which is to have the model conduct a critical reverse consistency check to eliminate false positive interference. This not only boosts the efficiency of subsequent steps but also ensures that only high-quality, genuine signals enter the downstream identification, improving identification accuracy.
3. Small Molecule Identification: The (c) Small Molecule Identification Agent is the analytical core of the system. It receives the predicted molecular formula and experimental spectral data from (b) and employs a dual-path analysis strategy for small molecule identification. This agent initially performs matching of the experimental spectrum against several spectral libraries, subsequently applying a cosine similarity algorithm to filter and retain the top-k library MS/MS spectra. If matching spectra are found, the agent will use the library-guided fragmentation analyzer for identification. This analyzer first assesses and preprocesses the data quality based on information such as retention time and signal-to-noise ratio to confirm the presence and accuracy of the molecular ion peak. Subsequently, the analyzer systematically compares the library and experimental spectra, identifies the key fragment ions, and seeks to link them to chemically sound fragmentation pathways originating from the precursor ion, offering detailed elucidation for each possible fragmentation route. Next, the analyzer performs a comprehensive evaluation by integrating multi-dimensional evidence, such as the relative abundance of fragment ions and isotopic distribution, to support or exclude a specific structural hypothesis. Ultimately, the analyzer synthesizes all preceding evidence to conclude whether the match “is the same compound,” “belongs to the same compound class,” or “does not belong to the same compound class”. If no matching spectrum is found, the agent employs the de novo fragmentation analyzer. The process of this analyzer is similar to the library-guided fragmentation analyzer, with the distinction that it will use structurally similar compounds for semi-unknown derivative analysis or conduct completely unknown compound analysis directly from the experimental data.
4. Results Aggregation and Reporting: Lastly, the (e) Reporting Agent functions as a central aggregator, gathering the MS data report from (a), the molecular formula prediction report from (b), and the small molecule identification report from (c). It then automatically generates a consolidated summary report for rapid review by researchers.

#### Instructions datasets

To improve the performance of the multi-agent system in small molecule identification, we constructed three supervised fine-tuning (SFT) instruction datasets: molecular formula prediction, false positive detection, and fragmentation analysis. Each instruction was collaboratively annotated by experts in small molecule identification and natural language processing. The small molecule identification experts were responsible for answering the instruction’s question, while the natural language processing experts handled prompt engineering^43^ and chain-of-thought (CoT)^44^ design. All instructions underwent a second round of manual cross-validation to ensure high data quality.

The molecular formula prediction dataset contains 1,000 instructions. The input for the instruction is a prompt composed of the ion mode, precursor ion m/z, and MS/MS fragment ion m/z. The output follows a designed CoT to perform analyses such as molecular weight calculation and fingerprint fragmentation based on the input information, thereby determining the molecular formula corresponding to the input experimental data. The false positive detection dataset contains 1,000 instructions. The input is a prompt composed of the compound’s elution retention time, ion mode, and MS/MS fragment m/z. The output performs reverse CoT reasoning based on the input information to determine if the input experimental data is a false positive. The fragmentation analysis dataset comprises 2,000 instructions, 1,000 of which are library-guided fragmentation analysis instructions. The input for this subset consists of both experimental spectra and the results of spectral-library matching. The output is composed of a CoT with four parts, which include data quality assessment and preprocessing, matching and identification of core fragmentation patterns and pathways, comprehensive assessment of multi-dimensional evidence, and final judgement and conclusion. Notably, when analyzing fragmentation pathways, the model utilizes a tree-of-thought (ToT)^45^ structure to explore various potential branches. The output will finally provide one of the following conclusions, that is “is the same compound,” “is the same class of compound,” or “is not the same class of compound. The other 1,000 de novo fragmentation analysis instructions are similar, with the difference being that they do not rely on library data for comparative analysis. Instead, they perform fragmentation analysis based only on the experimental data to directly derive the compound name corresponding to the molecular formula.

#### Foundation Model

Our work employs the ScienceOne^46^ scientific foundation model. The ScienceOne model has been deeply customized for scientific research and has demonstrated exceptional performance in core scientific domains. In this work, we use ScienceOne-14B as the foundation model for each agent. The model has a total of 14.8B parameters, contains 40 Transformer^47^ layers, a hidden layer size of 5120, a grouped-query attention^48^ mechanism with 40 attention heads and 8 key-value heads, and supports a context length of 32,768 tokens natively and 131,072 tokens using YaRN^49^.

#### Training

Both the molecular formula prediction model and the small molecule identification model were trained using the Adam optimizer, with hyperparameters β_1_ = 0.9 and β_2_ = 0.95. The learning rate was set to 5e-6, and was dynamically adjusted using a warmup and cosine annealing strategy to enhance model training stability. Weight decay was set to 0.1 to prevent overfitting during model training. A global random seed of 42 was used, and the training was conducted for 5 epochs.

To balance computational power and time costs, we applied techniques such as parallel training, gradient checkpointing, gradient accumulation strategy, fp16 precision, and Flash Attention 2^50^. The molecular formula prediction model underwent SFT on the molecular formula prediction and false positive detection instruction datasets. This training was performed on a cluster of 8 NVIDIA A100 80G GPUs, requiring 6 hours and 40 minutes per session, and was iterated 12 times. The small molecule identification model was fine-tuned using the fragmentation analysis instruction dataset. This process was also conducted on a cluster of 8 NVIDIA A100 80G GPUs, with each session lasting 5 hours, for a total of 10 training iterations. SFT used the negative log-likelihood (NLL) loss as the objective function. By minimizing the NLL of the model generating the target sequence on the training samples, it enables the model to learn to predict the correct next token given the input and previous outputs. The formula is as follows:

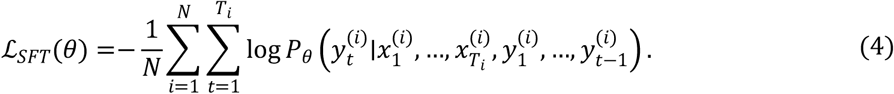

Where 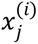 denotes input tokens (e.g., question or dialogue context) in the *i* -th example, whereas 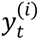 denotes the token at position *t* in the response. And, *P*_*θ*_( ·) denotes the probability of generating the token at position *t* given the input context and previously generated response tokens, computed by the model with parameters *θ*.

### Evaluation benchmark for molecular formula prediction

We selected a series of competitive baselines to compare the molecular formula prediction performance with our molecular formula prediction agent. The evaluation was conducted on MassSpecGym^51^, which is currently the largest public dataset containing high-quality annotated MS/MS spectra. Its quality assessment workflow ensures the reliability of molecular labels and metadata by filtering out noisy or corrupted spectra. Detailed descriptions of the baseline methods (MIST-CF^26^, FIDDLE^25^, BUDDY^21^, and SIRIUS^20^) are provided in Supplementary Note 1.

MassSpecGym includes three subsets: train, valid, and test. To ensure a fair evaluation and to verify the transferability of the methods, all methods were not trained on the train set but were instead evaluated directly on the test set. Statistics for the MassSpecGym test set are presented in Table 3.

We employed top-1 accuracy as the evaluation metric, where a prediction was considered correct only if the uniquely predicted molecular formula exactly matched the ground truth molecular formula; otherwise, it was deemed incorrect. The parameter settings for the baselines and our method are shown in Additional information Supplementary Table 1.

### Evaluation benchmark for small molecule identification

We initially performed LC-MS experiments on three complex TCM formulas: Buyang Huanwu decoction, Dengzhan Shengmai granule, and Zhuodu Huanwu decoction. The chemicals and reference standards used in this study are detailed in Supplementary Methods 1. Prior to analysis, samples were prepared following the specific extraction and concentration protocols described in Supplementary Methods 2. The LC-MS data acquisition was then conducted using the instrument parameters and gradient elution programs outlined in Supplementary Methods 3. Following this, we manually identified the small molecule constituents and verified them experimentally using the purchased reference standards. Table 4 presents the data statistics.

**Table 4.** MassSpecGym Test Set Data Statistics.

| Metric | Value |
| --- | --- |
| Number of records | 17,556 |
| Number of unique formulas | 2,777 |
| Molecular mass range | 213.99 ~ 996.31 |
| Unique elements in formulas | H, C, N, O, F, Si, P, S, Cl, Br, I |
| Unique adducts | [M+Na]<sup>+</sup>, [M+H]<sup>+</sup> |
| Unique instrument types | Orbitrap, QTOF, Unknown |

**Table 5.**
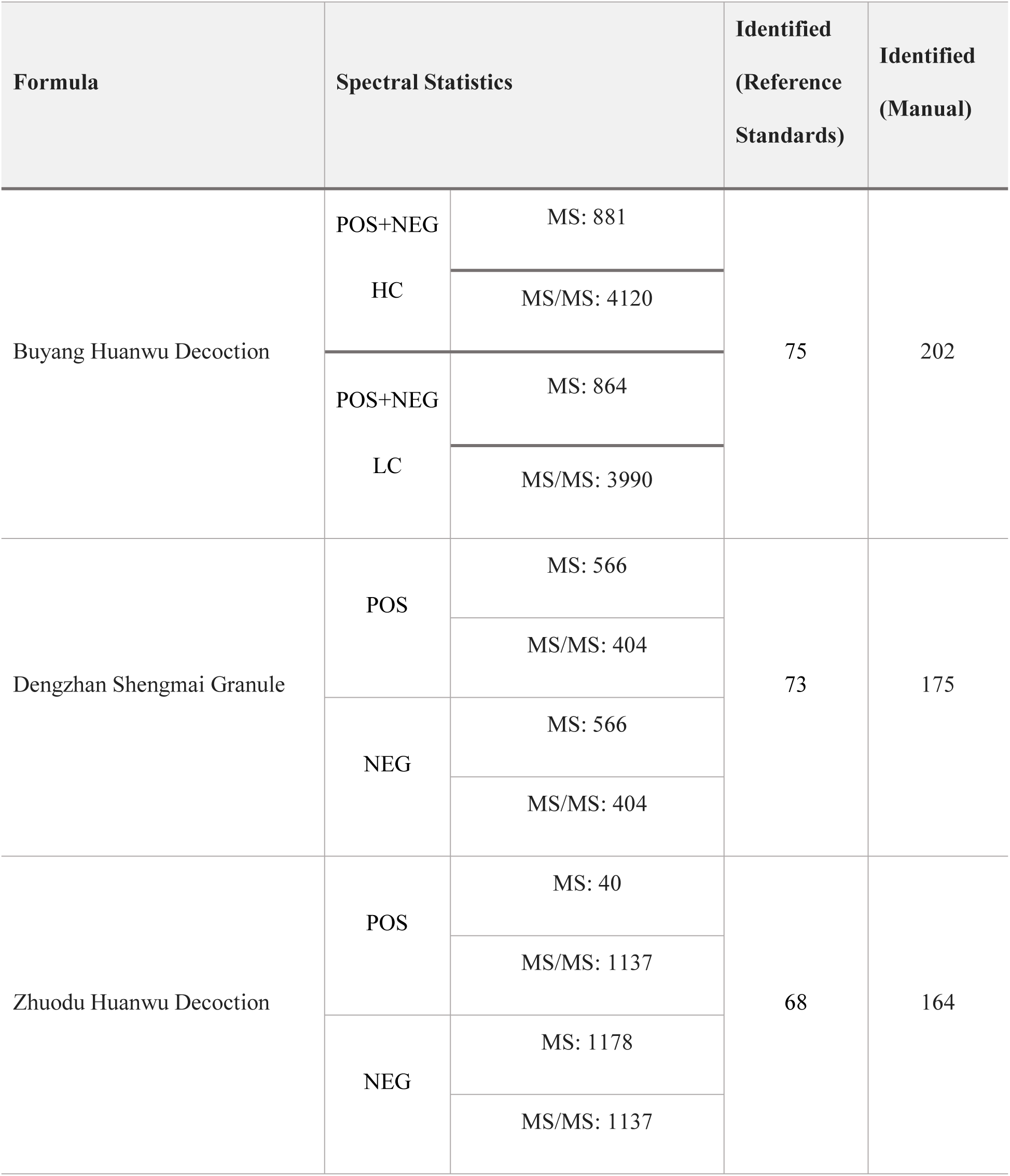
Statistical analysis of LC-MS experimental data.

Based on the aforementioned LC-MS experimental data, we used MS4MS to perform end-to-end small molecule identification. The molecular formula prediction model and small molecule identification model were deployed on 8× NVIDIA 4090 24G GPUs, with tasks executed serially. Subsequently, we submitted the identification results to a third-party organization for verification. Ultimately, we calculated the accuracy of the MS4MS identification results, comparing and analyzing them relative to the identification results.

